# Design, synthesis, and evaluation of a mitoxantrone probe (MXP) for biological studies

**DOI:** 10.1101/2023.04.11.536471

**Authors:** Savanna Wallin, Sarbjit Singh, Gloria E. O. Borgstahl, Amarnath Natarajan

## Abstract

Mitoxantrone (MX) is a robust chemotherapeutic with well-characterized applications in treating certain leukemias and advanced breast and prostate cancers. The canonical mechanism of action associated with MX is its ability to intercalate DNA and inhibit topoisomerase II, giving it the designation of a topoisomerase II poison. Years after FDA approval, investigations have unveiled novel protein-binding partners, such as methyl-CpG-binding domain protein (MBD2), PIM1 serine/threonine kinase, RAD52, and others that may contribute to the therapeutic profile of MX. Moreover, recent proteomic studies have revealed MX’s ability to modulate protein expression, illuminating the complex cellular interactions of MX. Although mechanistically relevant, the differential expression across the proteome does not address the direct interaction with potential binding partners. Identification and characterization of these MX-binding cellular partners will provide the molecular basis for the alternate mechanisms that influence MX’s cytotoxicity. Here, we describe the design and synthesis of a MX-biotin probe (MXP) and negative control (MXP-NC) that can be used to define MX’s cellular targets and expand our understanding of the proteome-wide profile for MX. In proof of concept studies, we used MXP to successfully isolate a recently identified protein-binding partner of MX, RAD52, in a cell lysate pulldown with streptavidin beads and western blotting.

**Graphical abstract (Draft):** 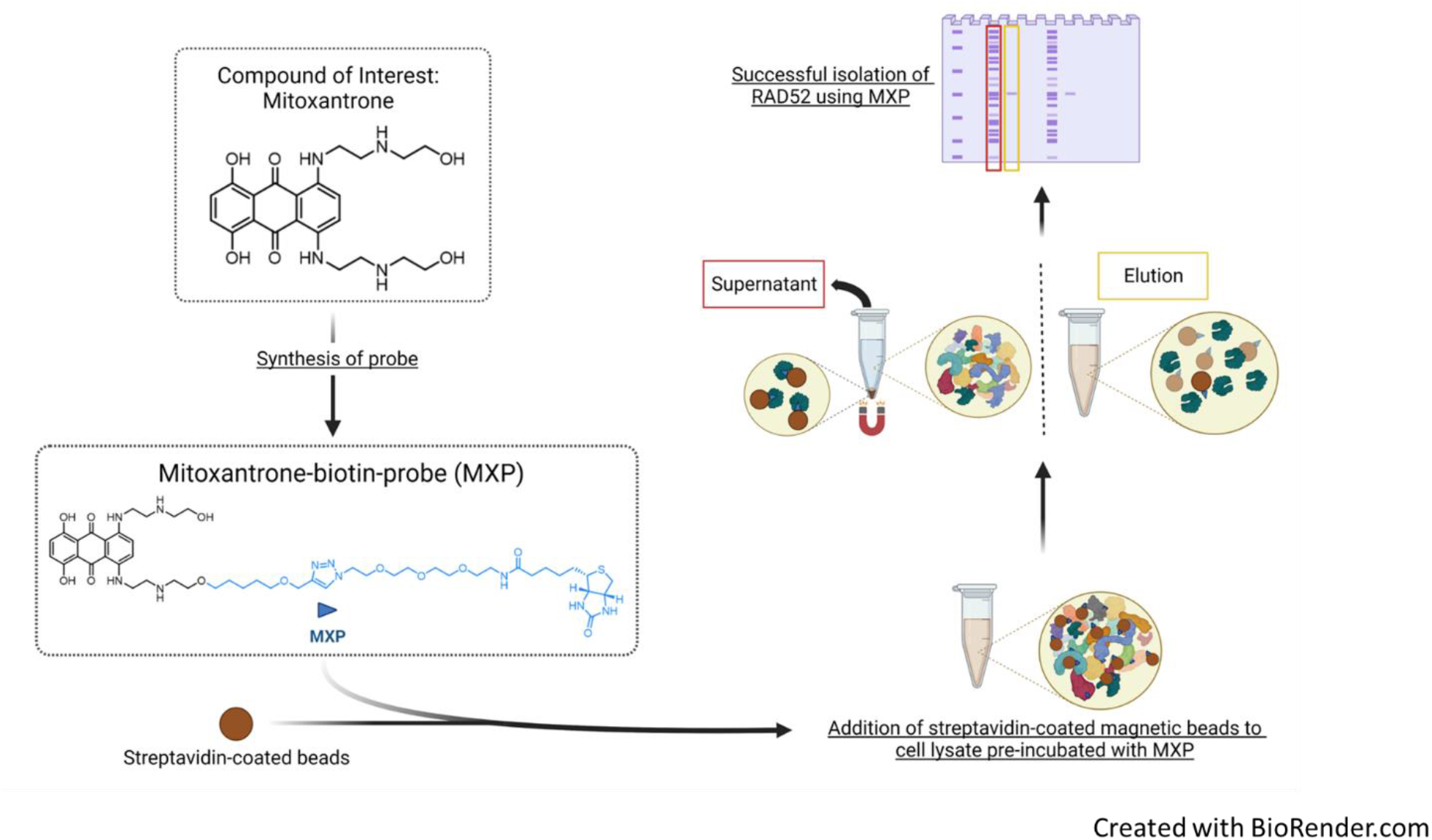

**Highlights:**

- An 8-step synthesis was used to generate a biotinylated-mitoxantrone probe (MXP).
- A pulldown of MXP demonstrated selectivity for RAD52, but not Replication Protein A.
- Western blot confirmed the identity of the isolated protein, RAD52.

---

Target identification and validation are key pillars of the drug discovery process.^1–3^ Biochemical probes facilitate the identification of drug targets or additional cellular binding partners that may contribute to off-target effects.^1, 4^ Such probes have driven target-based studies that profile drug interactions and elucidated the mechanism of action (MoA). These studies also guide future modification and optimization.^3, 5, 6^

In recent years, biotinylated-small molecule probes have had an immense impact on proteome-wide target profiling (Fig. 1). Examples include the biotinylated-resveratrol probe (Fig.1A) generated by Chen *et al*. that led to the identification of histone deacetylase I (HDAC1) as a cellular target of resveratrol.^7, 8^ Also, a biotinylated-triptolide probe (Fig. 1B) synthesized by Zhao *et al*. revealed peroxiredoxin I (Prx1) as a target of triptolide.^9^ Additional examples of biotinylated probes are included in Figure 1.^7–13^ These probe-based studies, and many others, are prime examples of using chemical biology tools to study complex interactions and interpret MoAs that are otherwise poorly understood.^5, 14, 15^

**Figure 1.**
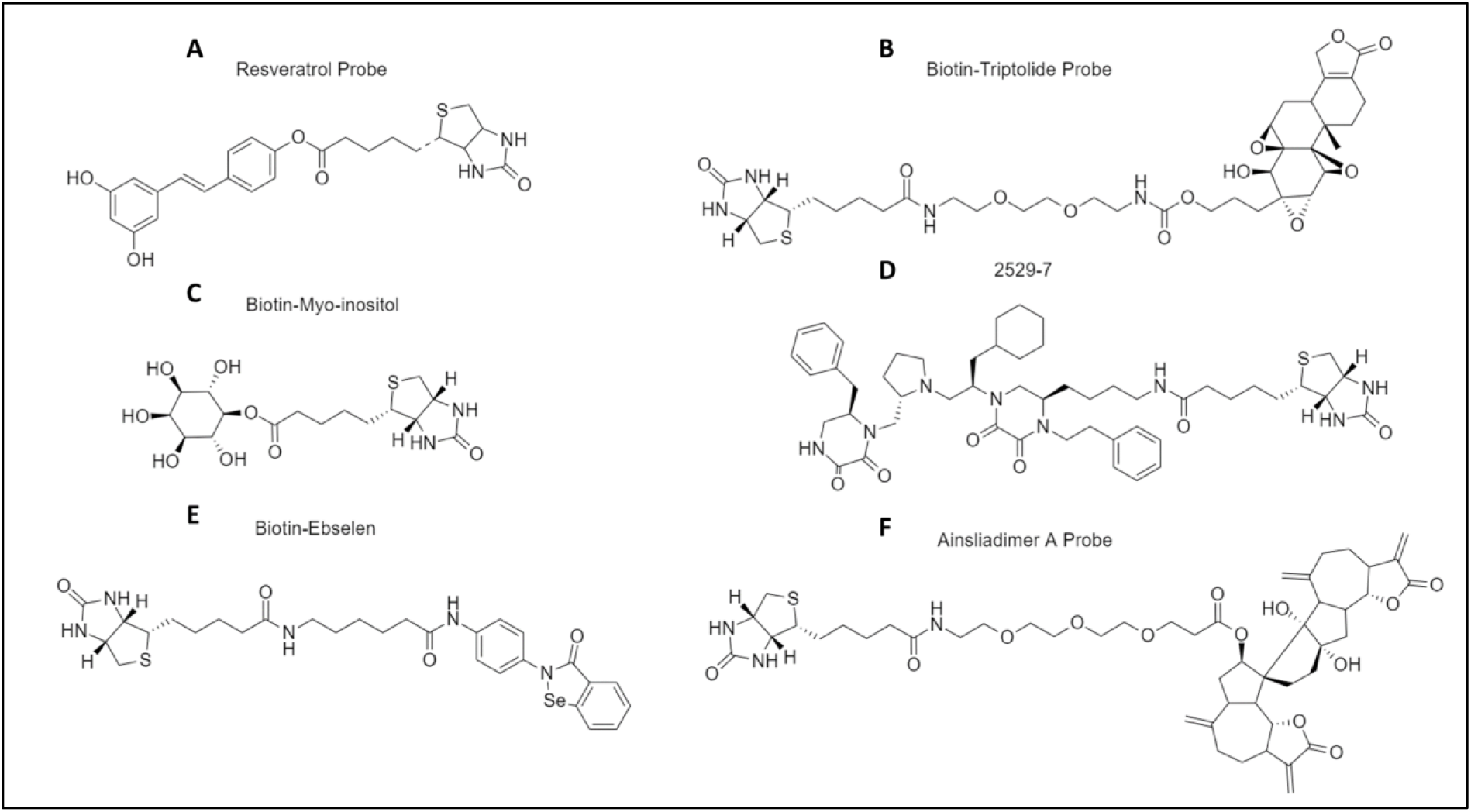
Examples of biotinylated small molecule probes used for mechanistic studies.

Mitoxantrone (MX) is a prominent anthracycline chemotherapeutic routinely used in oncology clinics. MX was originally approved in 1988 for the treatment of acute myeloid leukemia (AML) and has since shown promise in the treatment of other hematological malignancies, such as acute lymphoblastic leukemia (ALL) and acute nonlymphocytic leukemia (ANLL).^16^ Additionally, MX has shown efficacy in advanced breast and prostate cancers.^17^ The canonical MoA associated with MX is its capacity to intercalate DNA and stabilize the topoisomerase II cleavage complex.^16, 17^ MX was engineered to be a less cardiotoxic analog of doxorubicin that retains its therapeutic efficacy.^16, 17^ The divergence in cardiotoxicities between MX and doxorubicin is largely thought to be influenced by differences in membrane lipid peroxidation that generates reactive oxygen species to damage cardiac tissue.^16, 17^ Another factor is the selective inhibition of different isoforms of topoisomerase II (α and β).^16, 17^ Despite these findings, various biological assays continue to identify novel MX targets that may contribute to its anticancer activity.^18–27^

Although numerous studies have evaluated MX clinically and pharmacologically, its prominent DNA intercalation activity associated with MX overshadows additional potential cellular targets in the literature.^17^ Consequently, unique interactions of MX and their respective effects remain to be fully elucidated. In recent years, several research groups have begun unraveling additional targets of MX, demonstrating its promiscuous nature and underscoring the limit to our understanding of its ‘off-target’ effects.^17–27^ For example, our group previously identified MX as an inhibitor of the protein-protein interaction between RAD52 and Replication Protein A (RPA), a promising drug target for homologous recombination-deficient cancers.^27^ This discovery prompted our group to synthesize a probe that could be used to not only validate our previous findings but also advance the understanding of MX’s complex cellular interactions. It is clear that further proteomic-profiling studies are required to fully comprehend MX’s additional MoAs.

Proteomic analysis can reveal novel interactions of candidate compounds or existing drugs. Several strategies can be employed to determine the proteome-wide target profile of a given compound; these methods are generally compound-centric or mechanism-centric.^1^ Proteomic methods that are compound-centric typically modify the original compound to generate probes suitable for chemical biology studies such as target enrichment or fluorescent labeling of target proteins in cell extracts.^1, 2, 6, 14, 15, 28^ Click chemistry is often used to generate these probes, and thus, has majorly contributed to the understanding target profiles and MoAs.^5, 6^ The probe-isolated target proteins can be confirmed using western blotting or mass spectrometry. Although the biological probes are useful in deducing MoAs, they are challenging to generate since they must retain their bio-activity to be effective and reliable. In contrast, mechanism-centric proteomic methods analyze the differences in protein abundance, expression levels, post-translational modifications, and localization.^1, 3^ These methods provide insight into the cellular consequences associated with the compound but not necessarily defining the initiating event, *i.e*., target-binding.^1, 3^

Here, we focus on compound-centric chemical biology studies. Compound-centric studies reveal target protein profiles providing insights into differing molecular mechanisms and off-target effects.^1, 3^ This information can influence the continual optimization of the compound to increase efficacy and decrease inadvertent side effects.^3, 29^ For example, we previously reported the discovery of a spirocyclic analog (**19**) as an inhibitor of IKK*β*-mediated NF-*κ*B activation.^30, 31^ Subsequent studies using a biotinylated-analog 19 probe unveiled a proteome-wide target profile that identified over 330 proteins that are modified by analog 19.^32^ This led to the discovery of spirocyclic dimers (SpiDs) as promising anticancer agents that inhibited cancer cell growth, and induced apoptosis through activation of unfolded protein response.^32, 33^

To assess MX’s selectivity for RAD52 and to define the proteome-wide profile of MX, we synthesized a biotinylated-MX-probe (MXP). In a proof-of-concept, we demonstrated that MXP indeed selectively binds to recombinant human RAD52 but not RPA in *Escherichia coli* lysates that have an abundance of these two proteins. The clickable MXP can be used in biological studies to not only validate other targets of MXP but also define the proteome-wide target profile.

Scheme 1 depicts the strategy used to synthesize MXP and MXP-NC. In brief, S_N_2 displacement of one of the bromine atoms in 1,5-dibromopentane (**1**) by propargyl alcohol using sodium hydride as base yielded **2**. *N*-Boc protection of both nitrogen atoms of 2- ((2-aminoethyl)amino)ethan-1-ol (**3**) yielded the second linker fragment **4**. Under the Williamsons ether synthesis conditions, the S_N_2 displacement of the bromine atom in **2** by the hydroxyl group in **4** gave compound **5.** Deprotection of the Boc groups in **5** resulted in the key fragment with the alkyne tag **6**. Aluminum chloride catalyzed Friedel-Craft acylation between 4,7-difluoroisobenzofuran-1,3-dione (**7**) and hydroquinone (**8**) gave 1,4-difluoro-5,8-dihydroxyanthracene-9,10-dione (**9**) in 65% yield.^34^ The sequential ArSN2 displacement of the fluorine atoms in **9** by compounds **6** and **3** in DMF yielded **10** and **11**, respectively. The click reaction between alkyne tagged **11** and biotin-PEG3-azide resulted in the title compound, MXP as blue solid in 31% yield. A negative control, MXP-NC, was synthesized by S_N_2 displacement of bromine atom of 1-bromopentane (**12**) by propargyl alcohol followed by click reaction with biotin-PEG3-azide (Scheme 1). Additional information regarding methods and synthesis can be found in the suppl

**Scheme 1.**
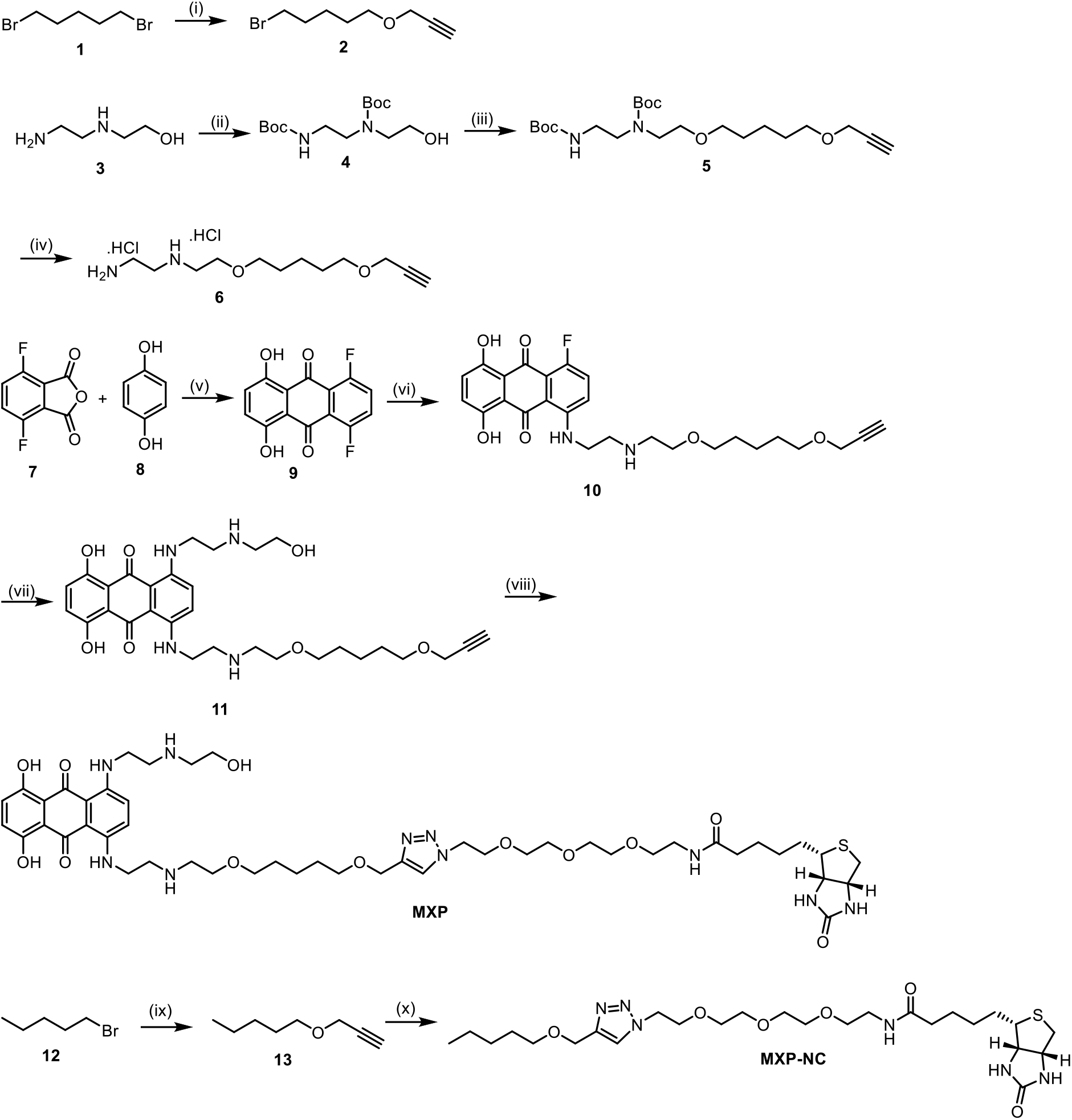
Reagents and conditions: (i) NaH, propargyl alcohol, DMF, 0°C to rt, 15 h; (ii) (BOC)_2_O, THF:EtOH (1:1), rt, 16 h; (iii) NaH, **2**, DMF, 0°C to rt, 15 h; (iv) 4M HCl in Dioxane, DCM, rt, 12 h; (v) AlCl_3_, NaCl, 200°C, 2 h; (vi) **6**, DIPEA, DMF, 50°C, 6 h; (vii) **3**, DMF, 50°C, 6 h; (viii) Biotin-PEG3-azide, Sodium abcorbate, TBTA, CuSO_4_.5H_2_O, DMSO, H_2_O, ^t^-BuOH, rt, 8 h; (ix) NaH, propargyl alcohol, DMF, 0°C to rt, 15 h; (x) Biotin-PEG3-azide, Sodium abcorbate, TBTA, CuSO_4_.5H_2_O, DMSO, H_2_O, ^*t*^-BuOH, rt, 8 h.

We previously reported that MX inhibited the RPA:RAD52 interaction using a high-throughput compatible FluorIA assay.^27^ We hypothesized that a MXP could be used to determine if MX binds to RAD52 or RPA using a pulldown study from *E. coli* lysates containing the two proteins. Figure 2 illustrates the pulldown of RAD52 using MXP in a RAD52-overexpressing *E. coli* cell lysate with magnetic streptavidin-coated beads. After overnight incubation of MXP and the cell lysate at room temperature, the magnetic beads were added to the sample for an additional 1-hour incubation. The supernatant fraction was stored after magnetic separation, and the beads were washed three times with lysis (wash) buffer. Following the 5-minute incubation with elution buffer (0.1 M Glycine, pH 2.0), the cellular components directly interacting with MXP were eluted from the probe, leading to an elution fraction enriched with binding partners. The presence of RAD52 was observed via SDS-PAGE gel stained with SYPRO™ Ruby dye (Figure 2A) and validated using an automated nanocapillary-based western blot system (Peggy Sue by Protein Simple) (Figure 2B). To evaluate whether MXP would isolate RPA, we included an *E. coli* cell lysate sample spiked with 0.06 mg/mL of purified RPA. Our SDS-PAGE results indicated that RAD52 was isolated in the lysate but was not immediately visible in the elution from the RPA-spiked lysate. Western blot was used for a more sensitive evaluation of our supernatant and elution samples.

**Figure 2.**
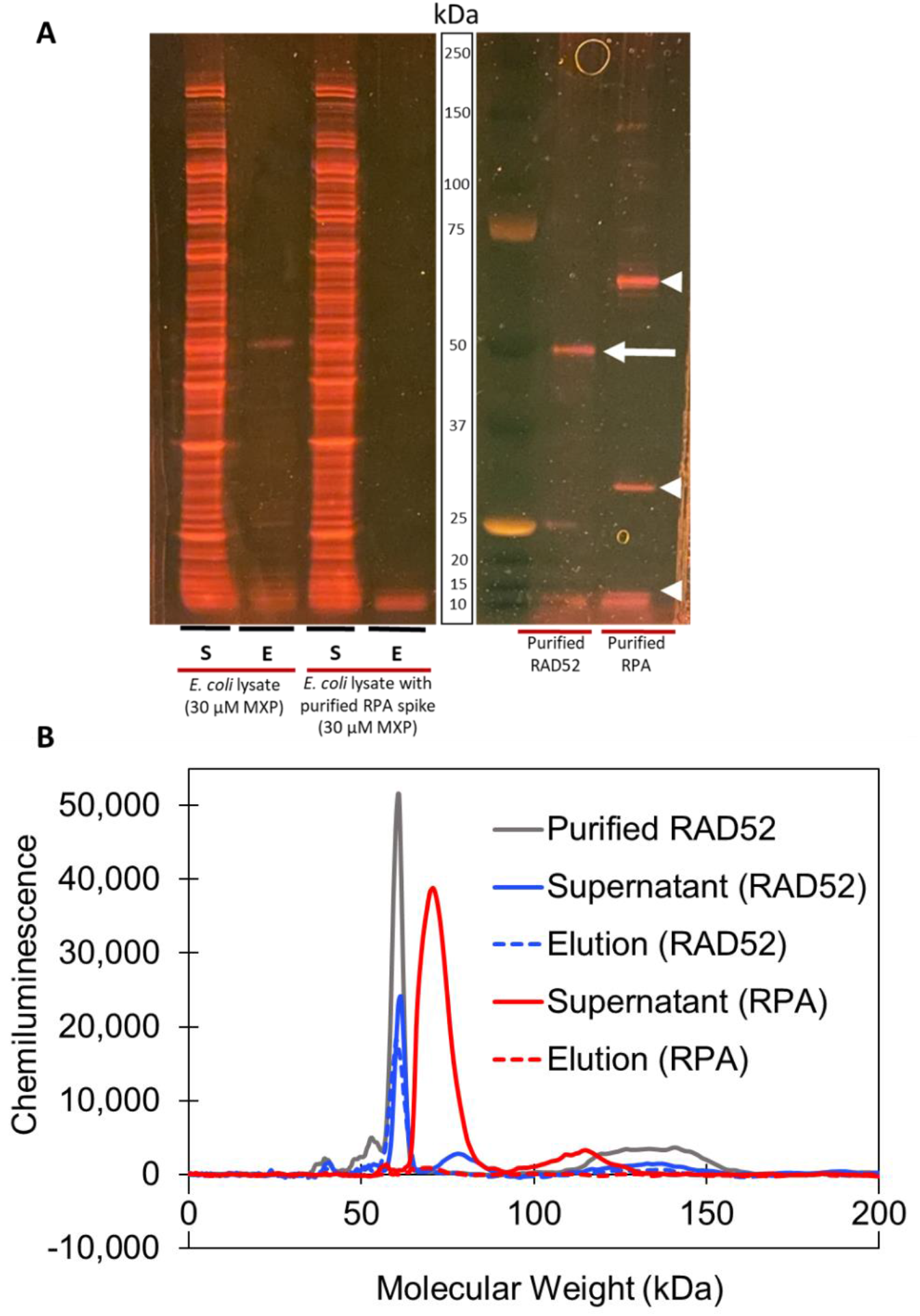
Isolating RAD52 via MXP pulldown in RAD52-overexpressing *E. coli* cell lysate. (A) SDS-PAGE gel stained with SYPRO™ Ruby dye showing supernatant (S) and elution (E) samples from a pulldown using MXP. A protein standard of purified RAD52 (white arrow) and purified RPA (white arrowheads) were included for reference. (B) Automated western blot results of the supernatant and elution using RAD52 antibody and RPA antibody for protein detection. Purified RPA was spiked into the *E. coli* lysate to determine if the MXP would interact with RPA or RAD52. As expected, RPA was only observed in the supernatant and not the elution, while RAD52, an identified protein-binding partner of MX, was detected in both the supernatant and the elution.

Peggy Sue western blots were performed following the Protein Simple manufacturer’s guidelines for the 12-230 size separation kit. Lysate samples were diluted to 0.04 mg/mL or 0.004 mg/mL as instructed. Primary antibodies against RAD52 and RPA (proteintech™ 28045-1-AP and Bethyl A300-241A) were used at a 1:200 dilution. Compass software (Protein Simple) was used to analyze the data. Further investigation using western blot found that RAD52 was in both the supernatant and elution of the RPA-spiked lysate samples. This indicated that the sensitivity of the SYPRO™ Ruby dye was too low to detect RAD52 in the RPA-spiked samples for the SDS-PAGE gel, but levels of RAD52 were detectable via a western blot.

Collectively, we have successfully synthesized a MXP that holds immense potential as a valuable tool for studying the chemical biology MX. The preliminary pulldown experiment demonstrated that MXP indeed interacts with cellular components and isolates a novel protein-binding partner. Moreover, it corroborates previous observations that MX interacts with RAD52, potentially implicating RAD52-inhibition as a promising complementary MoA in targeting homologous-recombination deficient cancers.^27^ Though MX has predominantly been thought of as a topoisomerase II poison, recent literature suggests the promiscuity of MX is understudied.^18–26^ Future studies using MXP may reveal additional binding partners and shed light on MX’s complex cellular interactions, enhancing our understanding of its proteome-wide target profile and additional modes of action that contribute to its therapeutic efficacy.

## Data availability

Data will be made available on request.

## Acknowledgements

This work was supported by funding from the CMDRP DOD OCRP (W81XWH-20-1-0816), Fred and Pamela Buffett NCI Cancer Center Support Grant (P30CA036727). SW was supported by student fellowships from Nebraska NASA EPSCoR Space Grant and NIH NCI training grant (T32CA009476). We thank Dr. Tadayoshi Bessho, Sneha Pandithar, and Dr. Mona Al-Mugotir for useful discussion.

## Author contributions

The manuscript was written through contributions of all authors. All authors have given approval to the final version of the manuscript.

Appendix A. Supplementary data

Supplementary data to this article can be found online.

