## Supplemental Data for "Design, synthesis, and evaluation of a mitoxantrone probe (MXP) for biological studies"

**General methods**

All reagents were purchased from commercial sources and were used without further purification. Flash chromatography was carried out on silica gel (200–400 mesh). Thin-layer chromatography (TLC) was run on pre-coated ANALTECH uniplate and observed under UV light at 254 nm and with basic potassium permanganate dip. Column chromatography was performed with silica gel (230-400 mesh, grade 60, Fisher Scientific, USA). ^1^H NMR (400 MHz) and ^13^C NMR (125 MHz) spectra were recorded in chloroform-d or DMSO-d6 on a Bruker-400 and Bruker-500 spectrometer (DMSO-d6 was 2.50 ppm for ^1^H and 39.55 ppm for ^13^C and CDCl_3_ was 7.26 ppm for 1H). Proton and carbon chemical shifts were reported in ppm relative to the signal from residual solvent proton and carbon. The data are presented as follows: chemical shift, multiplicity (s = singlet, d = doublet, t = triplet, q = quartet, p = pentet, m = multiplet and/or multiple resonances), coupling constant in hertz (Hz), and integration. HPLC was performed on a Waters Alliance 2695 system under the following conditions: column, Phenomenex Luna-2 RP-C18 (5 µm, 4.6 mm × 250 mm, 120 Å, Torrance, CA); solvent A, H_2_O containing 0.1% formic acid (FA); solvent B, CH3CN containing 0.1% FA; gradient, 10% B to 100% B over 15 min; injection volume, 10 µL; flow rate, 1 mL/min. The purified compound was further confirmed by HRMS analysis using Agilent 6230 time-of-flight LC/MS (LC/TOF) system.

Compounds **2**^1^, **4**^2^ and **9**^3^ were synthesized by using the reported procedures.

***tert*-butyl (2-((tert-butoxycarbonyl)amino)ethyl)(2-((5-(prop-2-yn-1-loxy)pentyl)oxy)ethyl)carbamate (5)**

Compound **4** (200 mg, 0.65 mmol) was dissolved in 1 mL of DMF. The mixture was stirred for few minutes at 0^o^C followed by portion wise addition of 60% NaH (26.28 mg, 0.65 mmol). The reaction mixture was further stirred for 30 minutes at same temperature followed by slow addition of **2** (134.7 mg, 0.65 mmol). The reaction mixture was stirred for additional 2 hours at 0^o^C and then at rt overnight. The reaction was quenched by adding ethyl acetate to the reaction mixture. The organic layer was washed subsequently with water and brine, respectively. The mixture was dried over Na_2_SO_4_, evaporated and purified by column chromatography using ethyl acetate/hexane (2/8) as eluting system; Yield = 38.7 % (109 mg); ^1^H NMR (400 MHz, CDCl_3_) δ 5.29 (br, 1H), 4.14 (d, *J* = 4.0 Hz, 2H), 3.53 (dd, m, 4H), 3.48 – 3.33 (m, 6H), 3.28 (m, 2H), 2.42 (t, *J* = 2.4 Hz, 1H), 1.70 – 1.53 (m, 4H), 1.55 – 1.36 (m, 20H); HRMS Calculated for C_22_H_41_N_2_O_6_^+^ m/z 429.2959, found mass = 429.2985

***N*1-(2-((5-(prop-2-yn-1-yloxy)pentyl)oxy)ethyl)ethane-1,2-diamine hydrogen chloride (6)**

To the solution of **5** (100 mg, 0.23 mmol) in DCM was added 4M HCl in dioxane (2 mL) at 0^o^C. The reaction mixture was stirred for 2 hours at 0^o^C and then at rt overnight. After completion of reaction, the solvent was removed under reduced pressure and crude product was washed with pentane and dried; Yield = 95.6% (71 mg); ^1^H NMR (400 MHz, MeOD) δ 4.15 (d, *J* = 2.4 Hz, 2H), 3.76 (dt, *J* = 17.8, 7.5 Hz, 2H), 3.56 (m, 4H), 3.51 – 3.25 (m, 6H), 2.85 (t, *J* = 2.4 Hz, 1H), 1.79 – 1.58 (m, 4H), 1.56 – 1.39 (m, 2H); HRMS Calculated for C_12_H_25_N_2_O_2_^+^ m/z 229.1911, found mass = 229.1907.

1- **fluoro-5,8-dihydroxy-4-((2-((2-((5-(prop-2-yn-1-yloxy)pentyl)oxy)ethyl)amino) ethyl)amino)anthracene-9,10-dione** (**10**)

To a solution of **9** (50 mg, 0.18 mmol) in DMF (1 mL) was added a mixture of **6** (54.5 mg, 0.18 mmol) and DIPEA (126 µL, 0.72 mmol) dissolved in 1 mL of DMF. The reaction mixture was stirred for 3 hours and then purified by HPLC; Yield = 13.9 mg (15.8%); ^1^H NMR (400 MHz, MeOD) δ 7.43 (t, *J* = 10.4 Hz, 1H), 7.38 – 7.29 (m, 1H), 7.23 (s, 2H), 4.11 (t, *J* = 4.9 Hz, 2H), 3.80 (t, *J* = 6.1 Hz, 2H), 3.76 – 3.65 (m, 2H), 3.53 (dt, *J* = 12.7, 6.4 Hz, 4H), 3.39 (t, *J* = 6.2 Hz, 2H), 3.33 (m, 2H), 2.82 (t, *J* = 2.4 Hz, 1H), 1.73 – 1.52 (m, 4H), 1.50 – 1.35 (m, 2H); ^13^C NMR (100 MHz, MeOD) δ 187.9, 185.9, 156.8, 156.2, 155.4, 148.4, 128.5, 127.5, 126.6, 126.3, 120.6, 119.4, 113.0, 112.9, 111.6, 74.2, 71.1, 69.4, 65.1, 57.3, 38.8. 28.9, 28.9, 22.4; HRMS Calculated for C_26_H_30_FN_2_O_6_^+^ m/z 485.2082, found mass = 485.2152.

**1,4-dihydroxy-5-((2-((2-hydroxyethyl)amino)ethyl)amino)-8-((2-((2-((5-(prop-2-yn-1-yloxy)pentyl)oxy)ethyl)amino)ethyl)amino)anthracene-9,10-dione (11**)

Compound **10** (8.4 mg, 0.017 mmol) was dissolved in 500 µL of DMF. To the mixture was added compound **3** (2.7 mg, 0.026 mmol) dissolved in 200 µL of DMF. The progress of reaction was monitored by HPLC and after completion of reaction after 6 hours, the product was purified by HPLC to yield **11** in 50.7 % yield (5 mg); ^1^H NMR (600 MHz, MeOD) δ 7.53 – 7.43 (m, 2H), 7.18 (s, 2H), 4.11 (t, *J* = 2.9 Hz, 2H), 3.96 – 3.80 (m, 6H), 3.80 – 3.68 (m, 2H), 3.66 – 3.46 (m, 4H), 3.45 – 3.38 (m, 3H), 3.25 (m, 2H), 2.88 – 2.75 (m, 1H), 1.72 – 1.53 (m, 4H), 1.50 – 1.37 (m, 2H); ^13^C NMR (150 MHz, MeOD) δ 189.2, 187.9, 156.9, 147.1, 129.7, 126.9, 125.0, 121.8, 119.2, 115.9, 111.7, 80.8, 75.6, 72.5, 70.8, 66.3, 58.7, 57.7, 51.0, 47.8, 39.9, 30.3, 30.2, 23.8; HRMS Calculated for C_30_H_41_N_4_O_7_^+^ m/z 569.2970, found mass = 569.3030.

***N*-(2-(2-(2-(2-(4-(((5-(2-((2-((5,8-dihydroxy-4-((2-((2-hydroxyethyl)amino)ethyl)amino)-9,10-dioxo-9,10-dihydroanthracen-1-yl)amino)ethyl)amino)ethoxy)pentyl)oxy)methyl)-1H-1,2,3-triazol-1-yl)ethoxy)ethoxy)ethoxy)ethyl)-5-((3aS,4S,6aR)-2-oxohexahydro-1H-thieno[3,4-d]imidazol-4-yl)pentanamide (MXP)**

To the stirred solution of biotin-PEG3-azide (4.72 mg, 0.010 mmol) and **11** (5.5 mg, 0.0096 mmol) in DMSO (1 mL) was added sodium ascorbate (6.70 mg, 0.0336 mmol) in H_2_O (300 µL) followed by TBTA (1.02 mg, 0.0019 mmol) in DMSO/H_2_O/tBuOH (3:1:1, 500 µL). To this solution was added CuSO_4_.5H_2_O (2.42 mg, 0.00096 mmol) in 80 µL of H_2_O and the reaction mixture was stirred for 8 hours at rt. The product was purified by reverse phase HPLC to give the title product in 30.6% yield (3 mg); HPLC purity 94%; ^1^H NMR (600 MHz, MeOD) δ 7.98 (s, 1H), 7.52 (s, 2H), 7.19 (s, 2H), 4.58 – 4.53 (m, 2H), 4.51 (s, 2H), 4.47 (dd, *J* = 7.8, 4.7 Hz, 1H), 4.27 (dd, *J* = 7.9, 4.4 Hz, 2H), 3.94 – 3.85 (m, 5H), 3.85 – 3.78 (m, 2H), 3.72 – 3.66 (m, 2H), 3.63-3.55 (m, 6H), 3.52 – 3.45 (m, 5H), 3.25 – 3.21 (m, 2H), 3.19 – 3.12 (m, 2H), 2.93 – 2.87 (m, 2H), 2.19 – 2.14 (m, 2H), 1.74 – 1.50 (m, 8H), 1.47 – 1.35 (m, 4H); ^13^C NMR (150 MHz, MeOD) δ 188.1, 176.1, 166.1, 157.0, 147.1, 145.8, 127.2, 125.8, 125.1, 115.9, 111.9, 72.4, 71.5, 71.4, 71.2, 70.6, 70.3, 66.3, 64.5, 63.3, 61.6, 57.7, 57.0, 51.4, 51.0, 41.1, 40.3, 40.0, 36.7, 30.3, 30.2, 29.7, 29.5, 26.9, 23.8; HRMS Calculated for C_48_H_72_N_10_NaO_12_S^+^ m/z 1035.4950, found mass = 1035.4929

**5-((3aS,4S,6aR)-2-oxohexahydro-1H-thieno[3,4-d]imidazol-4-yl)-N-(2-(2-(2-(2-(4-((pentyloxy)methyl)-1H-1,2,3-triazol-1-yl)ethoxy)ethoxy)ethoxy)ethyl)pentanamide (MXP-NC)**

**MXP-NC** was synthesized using the same procedure as used for the synthesis of **MXP-NC**; Yield = 43%; ^1^H NMR (400 MHz, MeOD) δ 4.60 (dd, *J* = 8.9, 3.6 Hz, 4H), 4.51 (dd, *J* = 7.8, 4.6 Hz, 1H), 4.32 (dd, *J* = 7.9, 4.4 Hz, 2H), 4.01 – 3.87 (m, 2H), 3.73 (s, 2H), 3.68 – 3.59 (m, 8H), 3.54 (m, 4H), 3.37 (m, 1H), 3.27 – 3.19 (m, 1H), 2.94 (dd, *J* = 12.7, 5.0 Hz, 1H), 2.72 (d, *J* = 12.7 Hz, 1H), 2.22 (t, *J* = 7.4 Hz, 2H), 1.86 – 1.52 (m, 6H), 1.53 – 1.41 (m, 2H), 1.35 (m, 4H), 0.93 (t, *J* = 7.1 Hz, 2H); HRMS Calculated for C_26_H_46_NaN_6_O_6_S^+^ m/z 593.3097, found mass = 593.3186

**^1^H NMR of MXP**

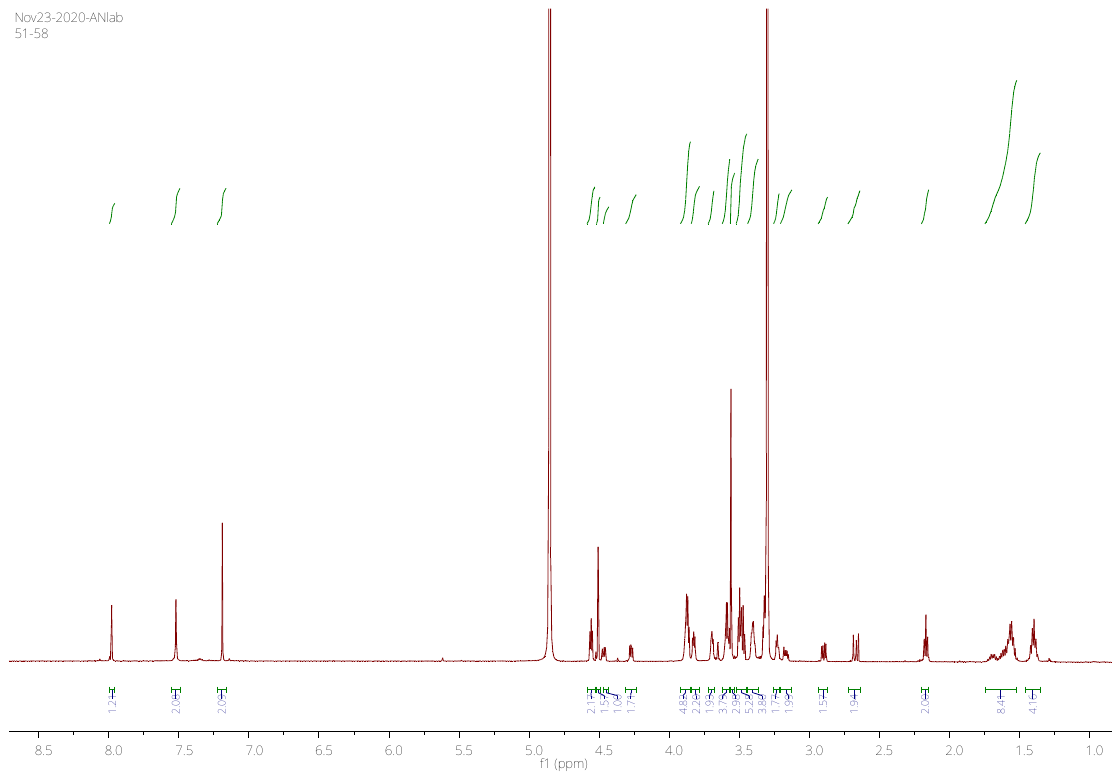

**^13^C NMR of MXP**

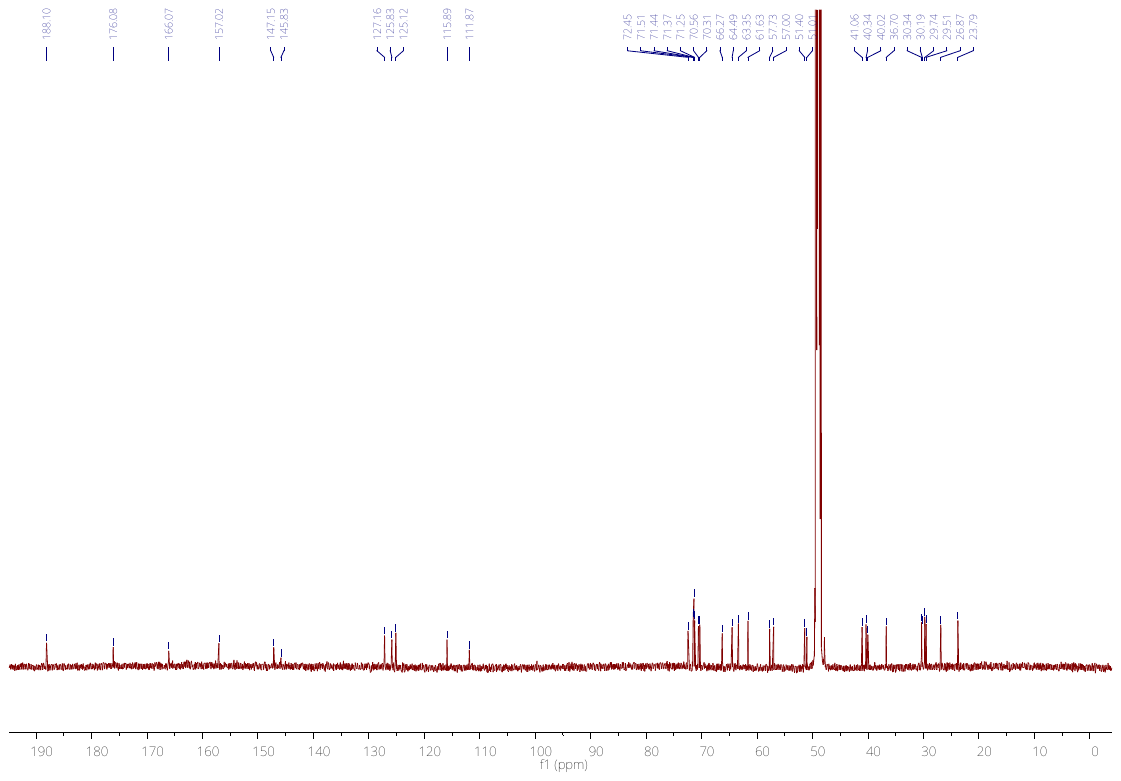

**HRMS spectra of MXP**

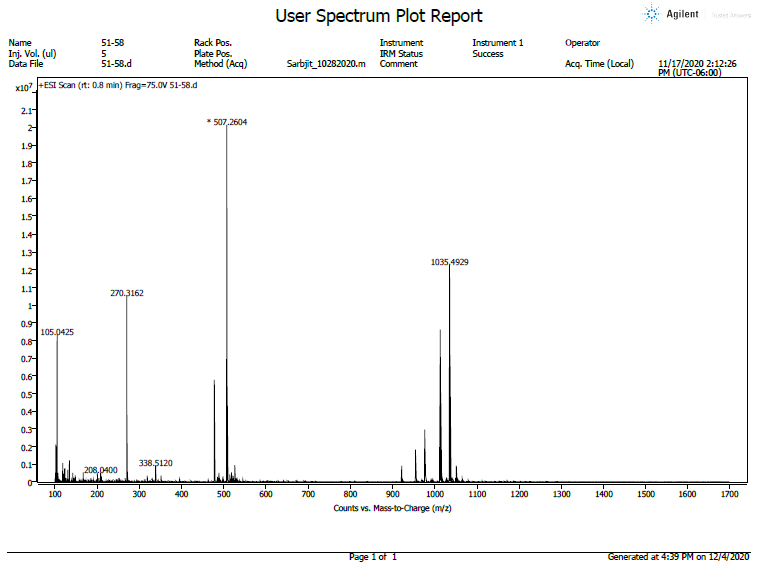
